# Directional swimming patterns in jellyfish aggregations

**DOI:** 10.1101/2024.03.08.584080

**Authors:** Dror Malul, Hadar Berman, Aviv Solodoch, Omri Tal, Noga Barak, Gur Mizrahi, Igal Berenshtein, Yaron Toledo, Tamar Lotan, Daniel Sher, Uri Shavit, Yoav Lehahn

## Abstract

Having a profound influence on marine and coastal environments worldwide, jellyfish hold significant scientific, economic, and public interest. The predictability of outbreaks and dispersion of jellyfish is limited by a fundamental gap in our understanding of their movement. Although there is evidence that jellyfish may actively affect their position, the role of active swimming in controlling jellyfish movement, and the characteristics of jellyfish swimming behavior, are not well understood. Consequently, jellyfish are often regarded as passively drifting or randomly moving organisms, both conceptually and in process studies. Here we show that the movement of jellyfish is controlled by distinctly directional swimming patterns, which are oriented against the direction of surface gravity waves. Taking a Lagrangian viewpoint from drone videos that allows the tracking of multiple adjacent jellyfish, and focusing the scyphozoan jellyfish *Rhopilema nomadica* as a model organism, we show that the behavior of individual jellyfish translates into a synchronized directional swimming of the aggregation as a whole. Numerical simulations show that this counter-wave swimming behavior results in biased correlated random-walk movement patterns that reduce the risk of stranding, thus providing jellyfish with an adaptive advantage critical to their survival. Our results emphasize the importance of active swimming in regulating jellyfish movement, and open the way for a more accurate representation in model studies, thus improving the predictability of jellyfish outbreaks and their dispersion, and contributing to our ability to mitigate their possible impact on coastal infrastructure and populations.

---

Jellyfish outbreaks exert a profound influence on marine and coastal environments worldwide, impacting ecosystem structure and functioning, biogeochemical cycles, and human well-being [1–5]. Despite their broad impact, our understanding and ability to predict jellyfish outbreaks and their subsequent dispersion, are characterized by a high level of uncertainty. A major source of this uncertainty is the lack of sufficient knowledge of the nature of jellyfish movement. Although there is evidence that jellyfish may actively affect their position [6–9], the role of active swimming in controlling jellyfish movement, and the environmental cues triggering and directing it, are not clear. Consequently, jellyfish are often regarded as passively drifting or randomly moving organisms, both conceptually [2, 10] and in environmental studies [11–13].

A natural framework to study jellyfish movement is rooted in the movement ecology paradigm, which attributes the temporal change in the position of an organism to four basic components, namely motion capacity, navigation capacity, internal state and external factors [14, 15]. In the case of jellyfish, motion capacity has been thoroughly addressed in a large number of laboratory experiments and numerical models, providing a mechanistic understanding of jellyfish swimming abilities, energetics, modes of swimming, turning mechanics, and the unique flow structures that are created [16–18]. Here we strive to achieve a fundamental understanding of the nature of jellyfish movement, by unveiling the interrelationships between the three other components of the motion ecology paradigm. As a conceptual framework, we center our analysis around the eminent risk of stranding, whose severity is intensified by the fact that jellyfish swarms are predominantly found in proximity to the coastline [19]. We hypothesize that due to the critical need of jellyfish to reduce the risk of stranding, both their internal state (i.e. the intrinsic factors affecting the motivation of jellyfish to move) and navigation capacity are linked to external factors associated with the threat of stranding, jointly acting to reduce this threat.

Evidence regarding the importance of directional movement in reducing jellyfish stranding was provided by Fossette et al. [9], who attributed swimming directionality to the strong tidal currents characterizing their study area in the Bay of Biscay. Here we elucidate the nature of jellyfish movement in the context of surface currents that are not dominated by a coastward component, such that current-oriented swimming would not necessarily reduce the risk of stranding. We focus on the Southeastern Mediterranean Sea, where the circulation is characterized by relatively weak tidal currents and strong along-shore currents [20, 21]. Our model organism is the scyphozoan jellyfish *Rhopilema nomadica*, which forms massive seasonal regional blooms [22, 23].

A useful tool in the study of jellyfish is aerial imaging from airplanes and drones, which provides synoptic non-intrusive observations of large numbers of adjacent individuals [24–28]. We expand the common utilization of aerial imaging in jellyfish research, and collect the required information using drone videos, which provide the time-varying perspective necessary for investigating organismal movement.

Drone data were collected in eight experiments during summertime jellyfish blooms in 2020-2022 (Fig S1) during the morning hours (6-10AM). In each experiment, a research vessel was directed to the heart of a spatially-dense jellyfish aggregation that was detected in real-time by an observer on a small aircraft flying simultaneously above. Upon arrival at the experiment site, a series of videos (mean duration 4:28 minutes *±* 1:27 sd ; Table S1) was recorded by a drone hovering at a fixed height, location, and orientation above the aggregation (Movie 1). The videos were analyzed in a Lagrangian framework, which tracked jellyfish along their trajectories (Fig. 1; Movie 2). At constant intervals along the trajectory of each jellyfish, we obtained the instantaneous swimming orientation (*α*), defined as the direction in which the jellyfish bell was pointing, based on the observed body positioning (top insert in Fig. 1). The drone data was also used to estimate the mean direction of surface currents [29] and of gravity waves [30]. In addition, the direction of long waves (i.e. swell) was extracted from the Copernicus Marine Environment Monitoring Service (CMEMS) Mediterranean Sea waves reanalysis .

**Fig. 1.**
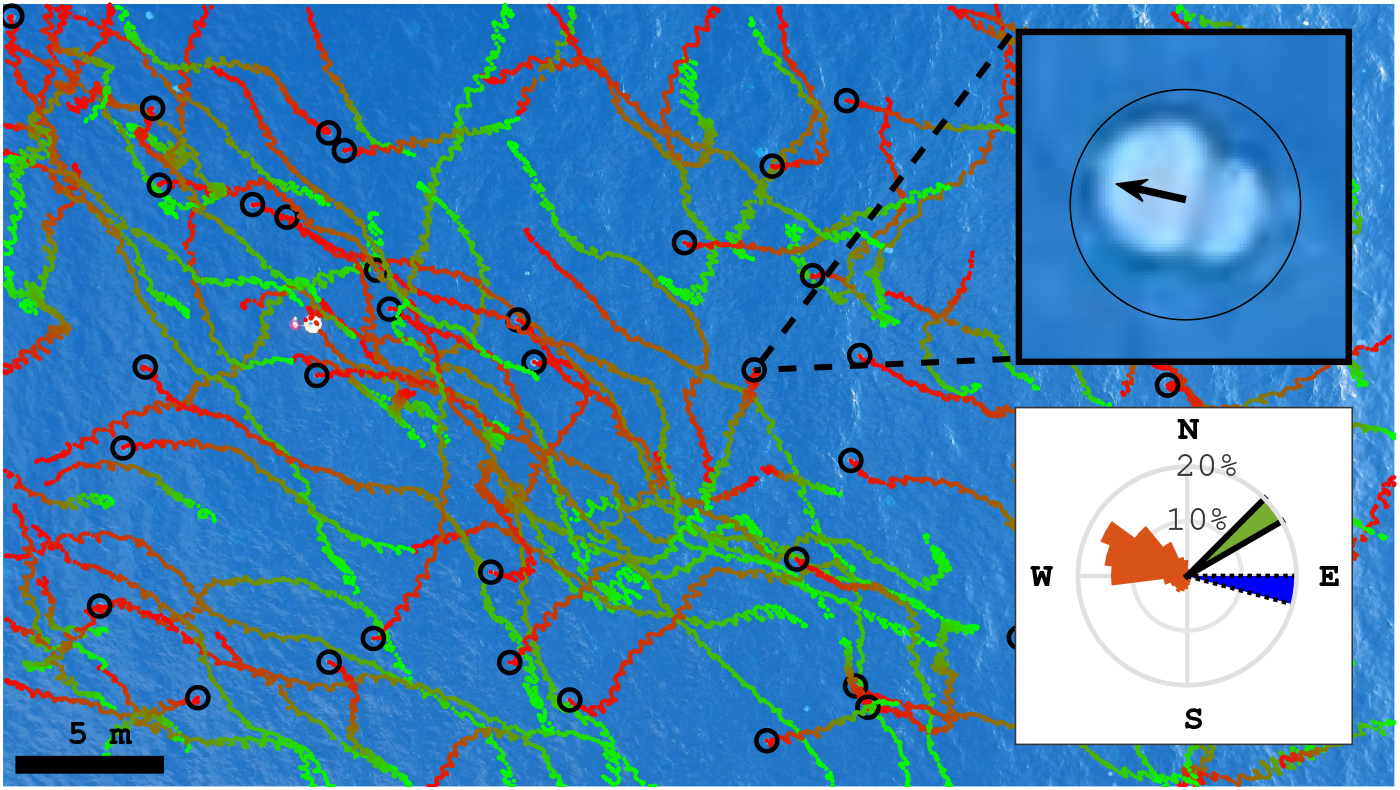
An exemplary drone-based Lagrangian view of the movement of aggregated jellyfish. The dotted lines show Lagrangian jellyfish trajectories extracted from a 5-minute video. The trajectories are overlaid on the last frame of the video, with the colors gradually changing from green to red during the trajectory. Black circles indicate locations of jellyfish in that frame. The upper insert shows an enlargement of a single frame, focusing on an individual jellyfish, with the black arrow indicating the swimming orientation, *α*. Lower insert shows the distribution of *α* for all instances measured in this video (2589 instances of 117 jellyfish in total, orange), and mean direction of surface gravity waves (blue, dotted edge) and currents (green, solid edge). The video was taken at 7:45 AM on July 6, 2020 (Fig. S1).

A key behavioral trait in aquatic locomotion is directional movement, defined as the tendency of an individual to move along a straight path [31, 32]. The observed jellyfish maintained a constant swimming orientation, with an average standard deviation of *α* along individual trajectories of 26.8 *±* 18.5^*°*^ (Fig. 2A). Out of the 4240 jellyfish examined, 4143 (*>* 97%) consistently exhibited statistically significant directional swimming (Rayleigh’s test *p <* 0.05 [33]).

**Fig. 2.**
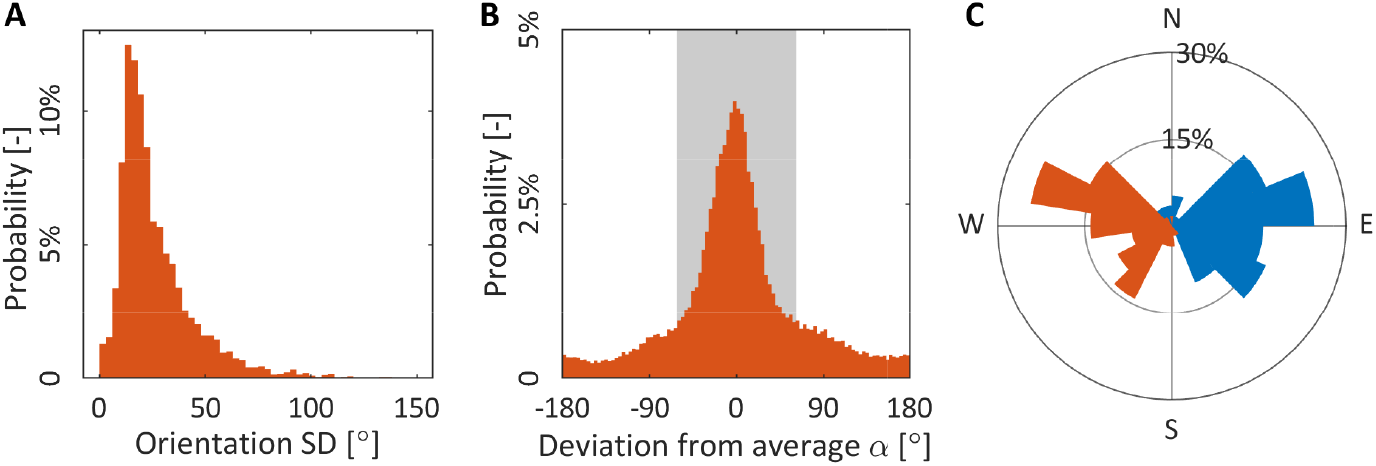
Characteristics of jellyfish swimming behavior. (A) The standard deviation of *α* along the trajectories of 4,240 jellyfish. The median standard deviation of *α* is 21^*°*^, which indicates that jellyfish maintained relatively straight paths. (B) The deviation of *α* from 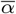 in all 90,429 instances of measurement (4,240 jellyfish), with the shaded area showing standard deviation. The narrow distribution indicated that aggregated jellyfish tend to swim in the same direction. (C) Mean direction of short surface gravity waves (blue) and 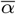 (orange) in each of the 57 movies examined.

Expanding the analysis, we tested swimming directionality at the scale of the jellyfish aggregation. For each video, we calculated the mean swimming orientation (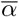) and found that the individual instantaneous orientations deviated from it by only *±*62*°* (shaded area in Fig. 2B). In agreement with this, aggregated jellyfish were found to collectively orient their swimming in the same direction (Rayleigh’s test *p <* 0.001). Notably, in all cases, 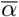 had a strong westward component, with a mean azimuth of 262 *±*44.7^*°*^ (north defined as 0^*°*^ and clockwise is the positive direction; Fig. 2C). In our study area, this westward orientation coincides with swimming away from the general direction of the coast (Fig. S1).

The directional nature of the swimming behavior indicates the use of an external cue [34]. Consistent with the hypothesized importance of stranding avoidance in modulating jellyfish movement, jellyfish swimming was distinctly oriented opposite to the direction of surface gravity waves, which in coastal areas provide a reliable indicator to the general direction of the shoreline [35], with 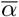 differing from the direction of short waves and long waves by 174 *±* 83^*°*^ and 155 50^*°*^, respectively (Fig. 2C and Table 1). Moreover, a statistical analysis revealed that 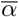 was significantly negatively correlated with the direction of long and short surface gravity waves (*p <* 0.001 ; circular-circular correlation; Table 1).

**Table 1.** Circular statistics between 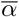 and possible environmental cues

| | $\rho$ | $p$ - value | Difference in direction<sup>1</sup> | $N$ | Source |
| --- | --- | --- | --- | --- | --- |
| Short surface waves | -0.658 | $p < 0.001$ | $174 \pm 83^\circ$ | 57 | Drone video |
| Long surface waves | -0.6731 | $p < 0.001$ | $155 \pm 50^\circ$ | 53 | [36] |
| Surface currents | 0.1363 | $p > 0.24$ | $115 \pm 77^\circ$ | 57 | Drone video |
| Sun azimuth | 0.4146 | $p < 0.01$ | $61 \pm 38^\circ$ | 53 | [37, 38] |
| Magnetic field delenation | 0.4755 | $p < 0.025$ | $103 \pm 40^\circ$ | 53 | [39] |

<sup>1</sup>Absolute values

Further investigation of the linkage between the different components of jellyfish movement was conducted through numerical modeling of jellyfish swimming behavior. We first reconstructed the observed jellyfish movement trajectories (e.g. Fig. 1). Jellyfish swimming speeds (*v*_*js*_) were taken from a normal distribution centered around 0.1 *ms*^*−*1^ with a standard deviation of 0.03 *ms*^*−*1^, based on a complementary analysis of the drone data used here [40]. The observed jellyfish trajectories were found to exhibit movement patterns that are best represented using a biased correlated random walk model (BCRW, [41]). To optimize the BCRW parameters, we employed a genetic algorithm, which yielded an angular diffusivity of 0.02 *s*^*−*1^, a preferred angle of 287^*°*^ and an average reorientation time to the preferred angle (*B*) of approximately 30 *s*. When subject to constant velocity current conditions, the simulated Lagrangian particle tracking produced trajectories similar to the drone-captured jellyfish trajectories (Fig. 3a).

**Fig. 3.**
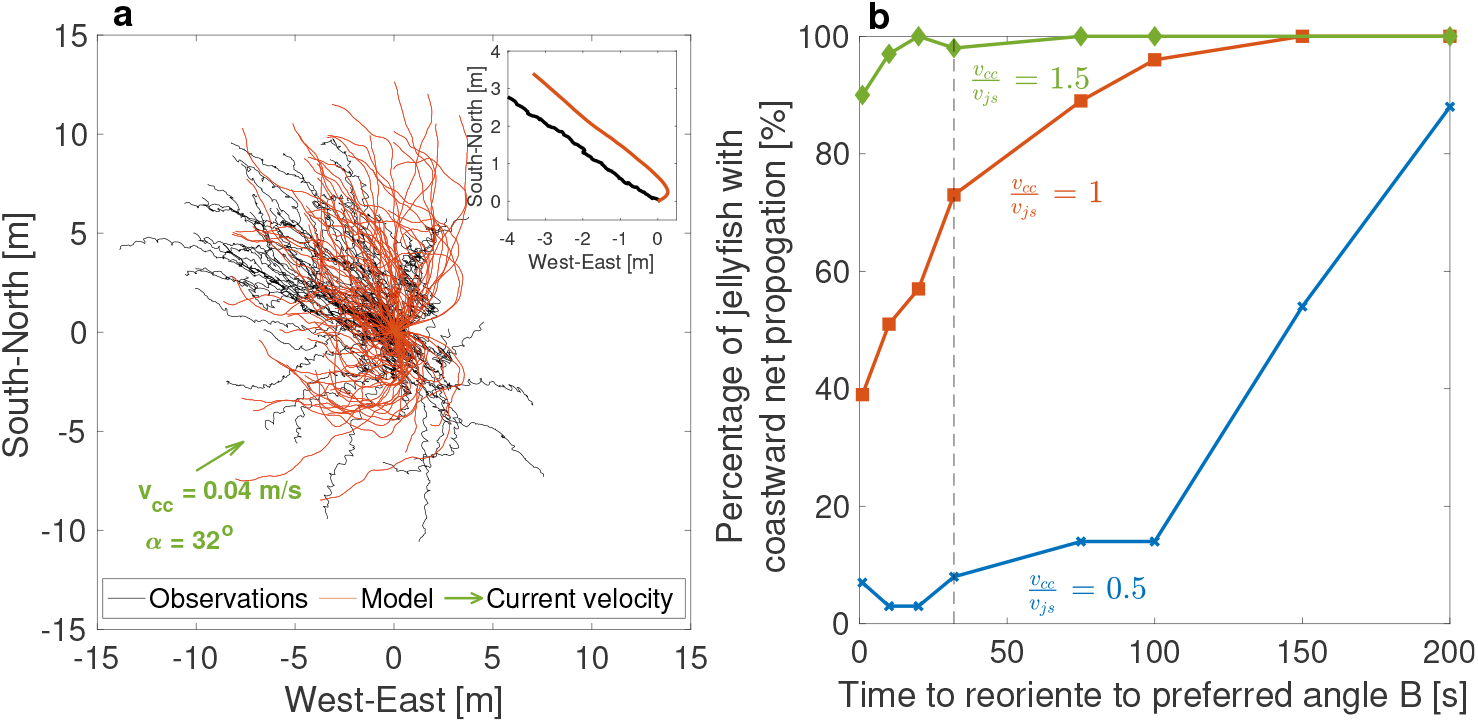
Numerical simulation of jellyfish swimming behavior and its impact on stranding. (a) Comparison between observed (black) and modeled (orange) jellyfish trajectories, for July 6, 2020. The model was run for 100 *s* under a constant current, *v*_*cc*_, of 0.04 *ms*^*−*1^, at an azimuth of 58^*°*^ (indicated by the green arrow in the lower left corner). The upper right insert displays the trajectories of center-of-mass of the observed (black) and modeled (red) aggregated jellyfish. (b) Percentage of jellyfish with net coastward propagation under varying model parameters *B* and 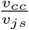 of 0.5 (blue, x symbol), 1 (red, square symbol) and 1.5 (green, diamond symbol). An increase in *B* manifest decrease in swimming directionality, with *B →* 0 representing fully directional swimming away from the coast, and 200 *s* represents simple random walk behavior). Vertical line marks the value of *B* found here.

To test the importance of directional swimming in reducing stranding risk, we compared the latter, defined here as the percentage of jellyfish whose net propagation was towards the coast, for varying levels of directionality (manifested by changes in *B*, going from 0 *s* for fully directional swimming away from the coast, to 200 *s* for simple random-walk behavior; Fig. 3b). The comparison was performed for coastward current speeds that are half, equal and 50% higher than the mean *v*_*js*_. These ratios are representative of the *R. nomadica* swimming speeds measured using the same drone data [40] and in laboratory experiments [42], and summertime surface currents measured in the region (Fig. S2). For the case of *v*_*js*_ equal to the coastward component of the current (*v*_*cc*_), the percentage of jellyfish propagating toward the coast went from 40% when the swimming was fully directional (i.e. *B →* 0*s*) to 100% when *B* approaches 150*s*. For the case of 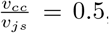, the percentage of jellyfish propagating toward the coast went from 7% when the swimming is fully directional to 88% when *B* increased to 200 *s*. When *v*_*cc*_ was very high compared with 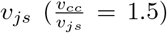, the directionality became negligible for jellyfish survival, as jellyfish swimming was too slow to counteract the flow (90% of jellyfish propagating toward the coast when *B →* 0*s*).

The westward (i.e., away from the coast) counter-wave swimming, and its role in reducing the risk of stranding, manifest a distinct relationship between the internal state and external forcing components of jellyfish movement. The opposite directionality and significant negative correlation between the wave and swimming directions suggest that an interrelationship also exists with the navigation capacity component, such that the jellyfish orient their swimming by perception of the surface gravity waves. This hypothesis is supported by the fact that in coastal areas, where jellyfish aggregations are commonly found [19], wave refraction causes the surface waves to be directed towards the coastline. This makes the applicability of a wave-perception mechanism for swimming away from the coast a universal feature. This is in contrast to other environmental cues that were previously found to be associated with jellyfish swimming directionality, such as the magnetic field [43], sun position [7], and current direction [9], which require a priori knowledge of the relative location of the coastline. In agreement with this, in our observations *α* was found to be less correlated with the sun azimuth and magnetic field inclination, and not significantly correlated with surface currents (circular-circular correlation *p >* 0.2) (Table 1). In addition, the actual process of beaching is typically driven by waves that, via Stokes drift, produce a principal mechanism for substantial cross-shore flows, as was recently shown in the case of oil pollution transport [44]. Therefore, efficient avoidance of stranding commonly requires counteracting the effect of waves, rather than that of the currents, which can only transport the jellyfish to the vicinity of the shoreline.

Evidence for animals orienting their swimming against surface waves is limited to a small number of animals [35, 45–48]. A wave-induced directional perception mechanism was identified in sea turtles, who were found to detect wave direction from the sequence of accelerations occurring within wave orbits below the water surface [49]. In jellyfish, while such a sensory mechanism has not been identified, counter-wave orientation was suggested as a possible explanation for the observed correlation between the direction of jellyfish swimming and that of surface wind [6]. As for the observations reported here, this explanation is supported by the fact that in the context of moving fluids, it is likely that any mechanoreceptor sensitive to fluid motion would not be sensitive to constant unidirectional flow, but rather to time-dependent components of the flow field. These may include shear flows, local turbulence, and orbital currents produced by surface waves [50, 51], as suggested here.

By providing a unique Lagrangian viewpoint on multiple adjacent jellyfish, our drone-based observations provide new insights into jellyfish swimming behavior and their resulting movement, in their natural environment. Focusing on aggregations of *R. nomadica*, we found that individual jellyfish consistently maintained a constant swimming direction, oriented against the surface gravity waves and away the shoreline. This behavior translates into synchronized directional swimming of the aggregation as a whole, which reduces the eminent risk of stranding, and provides jellyfish with an adaptive advantage critical to their survival. In addition to shedding light on jellyfish swimming behavior and its importance, our results open the way for a more accurate representation of jellyfish movement in model studies, thereby improving our ability to understand and predict dynamical, ecological, biogeochemical and societal aspects of jellyfish outbreaks.

## Methods

### Extraction of jellyfish trajectories and body orientation

The time-dependent locations and trajectories of individual jellyfish were obtained using the TrackMate plugin for Fiji [52, 53], which uses a blob detector based on a Laplacian of Gaussian (LoG) filter to detect circular object in the image, and a Linear Assignment Problem (LAP) tracker to generate trajectories. The plugin was designed for particle tracking in grayscale images and was used successfully in another study on drone-based jellyfish aggregations videos [54]. Videos were down-sampled from 30 frames per second (fps) to 2 fps (Movie 1), and converted to a HSV color-space. The value (V) layer of each HSV image was used as the grayscale image input to the plugin because it provided the highest visible contrast between the jellyfish and the background, yielding the best detection results. Movie 2 displays an example of a trajectory extraction result in the same video shown in Fig. 1. The trajectories of the jellyfish are marked with green to red points, with the change in color representing time. For reference, the white dots mark the trajectories of passive drifters (namely submerged plastic bags and bamboo plates [55]) without locomotive abilities.

Jellyfish body orientation was measured every 5 seconds using a manual method and a semi-automatic supervised method. In the manual method the orientation of individual jellyfish was manually marked (see example in the top insert in Fig. 1) using a custom MATLAB script. The supervised semi-automatic method also used a custom python script, which is based on fitting an ellipse engulfing the outline of the jellyfish. First, the image was cropped to a 50x50 pixels square centered around the location of the jellyfish in each frame. Next, we applied the Otsu method [56] on the red layer of the RGB image which resulted in a binary image, and fitted an ellipse to its edge. The long axis of the ellipse was defined to be the jellyfish body orientation, with the correct heading (out of the two possibilities) identified manually at least four times in each trajectory.

### Descriptive measures in circular statistics

To calculate the average and standard deviation of jellyfish orientation each instance of measurement was considered as a unit vector *r*_*i*_ = (cos *α*_*i*_, sin *α*_*i*_), where *α* is the angle and *i* is the index of the instance of measurement. The average unit vector therefore is calculated as an average of the unit vectors, 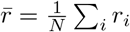, where *N* is the number of instances. The average angle 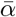 is computed using the 2-argument arctangent function,

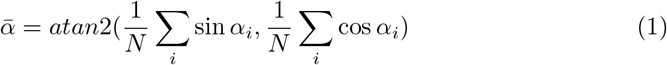

The standard deviation of an individual jellyfish trajectory (Fig. 2A) was calculated as,

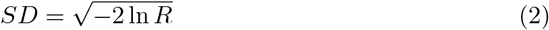

Where *R* is the absolute value of the average unit vector 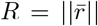. Means and standard deviations of angles were computed using a MATLAB toolbox for circular statistics [57].

### Statistical tests

Rayleigh’s test was used to test the directionality of 4240 individual jellyfish trajectories [32, 33] by testing the length of the resulting average vector length *R*_*j*_ for each jellyfish. The *P − value* is estimated as [58],

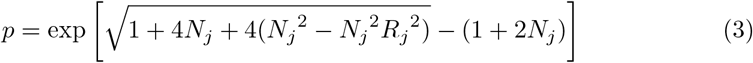

Where *N*_*j*_ is the number of orientation angles measured in a specific jellyfish trajectory. At a significance level lower than *p <* 0.05, orientations in a trajectory were considered non-uniformly distributed and thus directional

To measure the association between the mean direction of a swarm in a movie and environmental variables (listed in table 1) we used circular-circular correlation [33]. The correlation coefficient *ρ* is computed as,

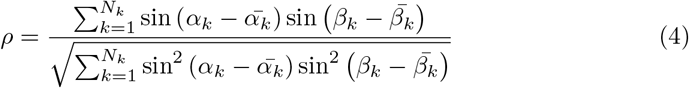

Where *N*_*k*_ is the number of videos used, *α*_*k*_ is the average orientation angle in a specific movie, 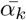 is a result of averaging *α*_*k*_ in all movies, *β*_*k*_ is any of the environmental variables listed in table 1 in a specific movie, and 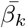 is the result of averaging the environmental variable in all movies. Statistical analyses were carried out in MATLAB using a toolbox for circular statistics [57].

### Drone-based measurements of surface waves and currents

Surface currents estimation from drone video was done following [29], using the CopterCurrents open-source MATLAB toolbox (https://github.com/RubenCarrascoAlvarez/CopterCurrents/releases/tag/1.0.01). The method infers the near-surface current based on its Doppler Shift of surface wave frequencies. The latter are estimated by fitting theoretical Doppler-shifted linear surface wave dispersion relation to the video gray-level Fourier spectrum. The assumption of linear-wave dispersion relation is justified here, as breaking waves (“white horses”) were very infrequent in the observed times and locations.

The bathymetric depths in the analyzed videos, a required input parameter in the estimation algorithm, were between 12-22 m according to the EMODnet bathymetric product [59]. Cross-checks with other bathymetric products yielded values within 10% differences. We have run the current analyses over wavelengths smaller than 15 meters, since wave frequencies depend weakly on bathymetric depth values as long as the values are similar or larger in magnitude than the wavelength. We have sverified that the estimated currents were indeed not sensitive to specification of bathymetric depth values differing by at least 10% from the EMODnet values.

Surface waves were estimated from the drone videos as well, via the method presented in [30]. In brief. the methodology involves computation of the space-time Fourier transform of the video (i.e., optical brightness spectrum), and inferring the sea surface wave spectrum via its relation to the optical brightness spectrum.

### Reanalysis swell estimates

To complement the short (wind-forced) waves spectrum estimated from the drone footage, we used long-wave (or swell) wave estimates from the CMEMS (Copernicus Marine Environment Monitoring Service) Mediterranean Sea waves reanalysis [36]. The reanalysis consists of a numerical simulation of sea surface wave physics, forced by best estimates of wind conditions, and constrained by daily observations of sea waves. The wave reanalysis product name in the CMEMS catalog is MEDSEA MULTIYEAR WAV 006 012. The reanalysis variable used here is “VMDR SW1”, i.e., the mean primary swell wave direction.

The reanalysis output files are available with hourly resolution, and we use the reanalysis conditions from its nearest available hour to each drone observation. We verified that reanalysis waves were similar in the previous or following adjacent hour in each case, justifying the use of this “nearest-neighbor” temporal interpolation approach. The reanalysis horizontal resolution is approximately 4 km. As such, its grid does not resolve Haifa Bay (where most of the drone footage was recorded). We therefore use the reanalysis wave conditions in its nearest grid cell to Haifa Bay. Given that long-waves are remotely generated, the swell inside the bay indeed enters from off-shore. The entering long waves can be modulated by refraction from the bathymetry. However, the spatial pattern of bathymetry within the sampling areas in the bay is relatively simple (not shown), and the observations were near the entrance to the bay. Hence refraction effects are expected to be minimal.

The swell mean period (specifically, the first inverse frequency moment, commonly written as T_1_) in the reanalysis for the relevant time and locations was 4.5-6 seconds, corresponding to wavelengths of *≈* 35-55 meters, in contrast with the range of wavelengths (*<* 15 meters) surveyed with the UAV analysis, as described above.

### Long term current measurements

Characteristics of the summertime currents in the region, which were used as an input for the numerical simulations, were derived from long-term continuous measurements performed by the Israel Oceanographic and Limnological Research (IOLR). The measurements were performed using an ADCP instrument located at the end of the Hadera pier, at distance of 2 *km* from the coast and at 27 *m* water depth. To represent conditions that are representative of our observations we used measurements taken between 6AM and 10AM, during June and July of the years 2020 - 2022. We used data from the upper cell of the ADCP, which corresponds to water depth of 4 *m*. The mean current speed was 18.5 *±* 10.9 *cms*^*−*1^. The current had a eastward (i.e. towards the coast) component during 69% of the time, with a mean zonal velocity of 4.3 *±* 7.3 *cms*^*−*1^ eastward.

### Numerical simulations

This section provides details on the Biased Correlated Random Walk (BCRW) model used for simulating jellyfish swimming, the observational data utilized for model optimization and validation and the Genetic Algorithm (GA) employed for optimization, and its results. Additionally, it explains the incorporation of passive drift into the active swimming model and provides details regarding the stranding simulations.

#### Biased Correlated Random Walk model

We used a BCRW as a simple 2D numerical model to characterize jellyfish active swimming [60]. This model is similar to the classic random walk model, however it incorporates a biased swimming direction where each step is correlated with the previous one. It has been previously used to characterize the movement of motile algal cells that are swimming in a preferred direction by gravitaxis and phototaxis [61], and also applied to the movement of larger animals such as ants [62], fish larvae [63] and seals [64]. We note that vertical swimming is neglected in this model for simplification, although we recognize that it may have a significant impact on jellyfish trajectories.

In the model, the equations governing jellyfish swimming trajectories are as follows:

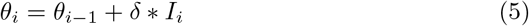

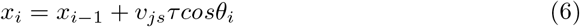

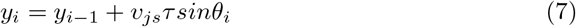

where *i* is the time step, (*x, y*) is the two-dimensional location of the jellyfish, *v*_*js*_ is the jellyfish swimming velocity, *τ* is the time step, *δ* is the angle step, *θ* is the angle of rotation and *I*_*i*_ is the direction of turning with values of (-1,1,0) determined by the probability to turn (see next paragraph). All fixed and free parameters and their values are detailed in Table 2. The jellyfish are initialized from the origin ((*x*_0_, *y*_0_) = 0) and *θ*_0_ is initialized randomly between 0-360^*°*^.

**Table 2.**
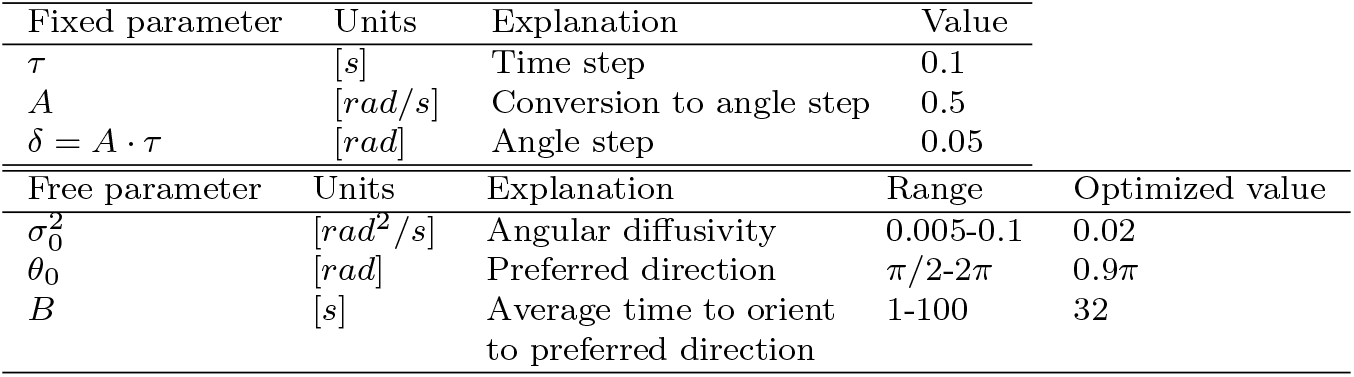
Parameters of the BCRW model. The model consists of two fixed parameters (one derived by the other two) and three free parameters optimized by the GA (6).

The model incorporates three free parameters that characterize the directionality of jellyfish active swimming: angular diffusivity (*σ*), preferred direction (*θ*) and the average time required to reorient to the preferred direction (*B*). The combination of these parameters determine the direction and directionality of jellyfish swimming through a probability to turn at each time step expressed as:

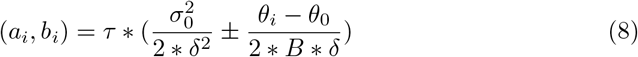

where *a* and *b* are the probability to turn clockwise and anticlockwise. The probability not to turn is, therefore 1 *− a − b*. According to the calculated probability the variable *I*_*i*_ assumes the values of (-1,1,0). Correlation with the previous step is evident in Eq. 6, as *θ* is incrementally adjusted in small steps at each time step, rather than being chosen randomly.

#### Model parameter optimization

The BCRW model parameters were optimized using a GA based on the trajectories obtained from drone footage taken in four days of field expeditions (24-25/6/2020, 6/7/2020 and 19/7/2020). Jellyfish swimming velocity was taken from a normal distribution with an average of 0.1 *±* 0.03 *ms*^*−*1^, as was found using the same drone data [40].

GAs have been used in marine science by ecosystem models such as NPZ and NPZD [65–67]. The optimization process aimed to minimize the cost function, the Mean Squared Error (MSE) of the center of mass of the trajectories. It’s worth noting that we also experimented with a Kullback-Leibler (KL) divergence cost function; however, it did not yield results as favorable as the MSE. The cost function was thus defined as follows:

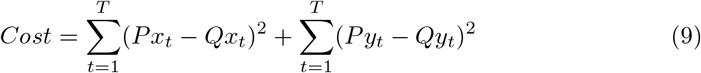

where *Q* represents the center of mass of 100 reconstructed time-dependent jellyfish trajectories, and *P* represents the center of mass of 100 modeled jellyfish swimming trajectories. The coordinates *x* and *y* correspond to the east-west and north-south axes, respectively (e.g., *Px* signifies the center of mass of modeled jellyfish trajectories along the x-axis). *t* denotes the time step, with a maximum of *T* = 100*s*.

The Genetic Algorithm (GA) ran for a total of 100 generations, with a precision of 10 bits (allowing for 1024 discrete values) within a specified range (see table 2). The population size was set at 10, and the probabilities for crossover and mutation were determined as 0.5 and 0.01, respectively.

The comparison between the model and the reconstructed 100 jellyfish trajectories after 100 seconds is depicted in figure S3a. In this figure, it’s evident that the model successfully captures the primary characteristics of jellyfish swimming, including the preferred direction, the trajectory length, and their extent in the north-south direction (y-axis). However, the model was unable to reconstruct the few trajectories where the jellyfish swim to the east (positive x-axis).

Figure S3b shows the center of mass of the reconstructed trajectories by date (those used and unused by the optimization procedure) and the modelled trajectories. As can be seen, the model fits in between the observations, both resembling the 24/6 and 6/7 which the model was optimized for, and the 25/6 and 19/7 used as validation. Overall, all swimming trajectories seem to have a similar swimming pattern, with a preferred direction in the northwest.

#### Modelling jellyfish trajectories under constant current

We further implemented the active swimming model into a Lagrangian particle tracking framework (OceanParcels, [68]). This framework adds a passive drift component of the flow to simulate the full jellyfish trajectories, containing both active swimming and passive drift. We added a constant current in specific directions as calculated by [40] during the same time of the observations. The results of the simulations for four different observations and velocities (19/7/2020 is split into two due to different velocity regimes that day) are depicted in Figure S4. As can be seen in figure S4, the model delivers favorable results for different flow regimes. As can be seen, the trajectories do not necessarily follow the passive drift, and in certain cases (6/7 and 19/7a) the full trajectories result in a propagation in the opposite direction (in the x-axis) to that of the current.

#### Stranding simulations

The purpose of the stranding simulations was to evaluate how the jellyfish’s resistance to stranding was affected by their directional swimming behavior. For simplicity, in these simulations the preferred direction was directed to the east (instead of northeast as found by the optimization). Assuming that jellyfish swim away from shore, the simplicity indicates that the shore lies on the east. Consequently, jellyfish that propagate to the west under the influence of the constant current would eventually become stranded.

The simulations were performed with 100 jellyfish and run for *∼* 10 hours. It is worth noting that a sensitivity test involving an increased number of jellyfish (1000) did not yield significant differences. In these simulations, *B* varied in the range of 1-200s, where the directionality increased as *B* decreased. The current velocity (*v*_*cc*_) was set at three different values: 5, 10, and 15 *cms*^*−*1^. The percentage of stranded jellyfish was estimated as the proportion of jellyfish that advanced a distance of 100 *m* or more towards the coast. This calculation assumed that jellyfish propagating 100m to the east would continue in that direction. To gain insight into the jellyfish behavior in these simulations, Figure S5 shows the jellyfish trajectories under varying values of *B* and *v*_*cc*_.

## Supplementary information

Supplementary Information is available for this paper

## Notes

### Competing Interest Statement

The authors have declared no competing interest.

