## Supplementary Information for "Directional swimming patterns in jellyfish aggregations"

**Table S1** A list of the drone footage durations and locations.

| Serial number | Date | Flight | Video | Latitude | Longitude | Duration [s] |
|---|---|---|---|---|---|---|
| 1 | 24/06/2020 | 1 | 2 | 32.85 | 34.95 | 409 |
| 2 | 24/06/2020 | 1 | 3 | 32.85 | 34.95 | 409 |
| 3 | 24/06/2020 | 1 | 4 | 32.85 | 34.95 | 205 |
| 4 | 24/06/2020 | 2 | 2 | 32.85 | 34.95 | 409 |
| 5 | 24/06/2020 | 2 | 3 | 32.85 | 34.95 | 409 |
| 6 | 24/06/2020 | 2 | 4 | 32.85 | 34.95 | 173 |
| 7 | 24/06/2020 | 3 | 1 | 32.85 | 34.95 | 242 |
| 8 | 24/06/2020 | 3 | 2 | 32.85 | 34.95 | 409 |
| 9 | 24/06/2020 | 3 | 4 | 32.85 | 34.95 | 198 |
| 10 | 24/06/2020 | 4 | 2 | 32.85 | 34.95 | 409 |
| 11 | 24/06/2020 | 4 | 3 | 32.85 | 34.95 | 409 |
| 12 | 24/06/2020 | 4 | 4 | 32.85 | 34.95 | 173 |
| 13 | 25/06/2020 | 3 | 3 | 32.84 | 34.99 | 353 |
| 14 | 25/06/2020 | 3 | 4 | 32.84 | 34.99 | 121 |
| 15 | 25/06/2020 | 4 | 1 | 32.84 | 34.99 | 352 |
| 16 | 25/06/2020 | 4 | 2 | 32.84 | 34.99 | 344 |
| 17 | 25/06/2020 | 4 | 4 | 32.84 | 34.99 | 301 |
| 18 | 06/07/2020 | 1 | 2 | 32.84 | 34.99 | 235 |
| 19 | 06/07/2020 | 1 | 3 | 32.84 | 34.99 | 308 |
| 20 | 06/07/2020 | 2 | 1 | 32.84 | 34.99 | 314 |
| 21 | 06/07/2020 | 2 | 2 | 32.84 | 34.99 | 327 |
| 22 | 06/07/2020 | 2 | 4 | 32.84 | 34.99 | 225 |
| 23 | 06/07/2020 | 3 | 1 | 32.84 | 34.99 | 231 |
| 24 | 06/07/2020 | 3 | 2 | 32.84 | 34.99 | 327 |

| | | | | | | |
|---|---|---|---|---|---|---|
| 25 | 06/07/2020 | 3 | 4 | 32.84 | 34.99 | 327 |
| 26 | 06/07/2020 | 4 | 1 | 32.84 | 34.99 | 294 |
| 27 | 06/07/2020 | 4 | 2 | 32.84 | 34.99 | 327 |
| 28 | 06/07/2020 | 4 | 4 | 32.84 | 34.99 | 305 |
| 29 | 06/07/2020 | 5 | 1 | 32.84 | 34.99 | 160 |
| 30 | 06/07/2020 | 5 | 2 | 32.84 | 34.99 | 198 |
| 31 | 06/07/2020 | 5 | 3 | 32.84 | 34.99 | 240 |
| 32 | 06/07/2020 | 5 | 4 | 32.84 | 34.99 | 196 |
| 33 | 19/07/2020 | 1 | 1 | 32.85 | 34.97 | 310 |
| 34 | 19/07/2020 | 1 | 2 | 32.85 | 34.97 | 273 |
| 35 | 19/07/2020 | 1 | 3 | 32.85 | 34.97 | 327 |
| 36 | 19/07/2020 | 2 | 6 | 32.84 | 34.97 | 188 |
| 37 | 19/07/2020 | 2 | 7 | 32.84 | 34.97 | 286 |
| 38 | 19/07/2020 | 2 | 8 | 32.84 | 34.97 | 272 |
| 39 | 19/07/2020 | 3 | 11 | 32.85 | 34.95 | 237 |
| 40 | 19/07/2020 | 3 | 12 | 32.85 | 34.95 | 241 |
| 41 | 19/07/2020 | 3 | 13 | 32.85 | 34.95 | 280 |
| 42 | 19/07/2020 | 3 | 15 | 32.85 | 34.95 | 290 |
| 43 | 19/07/2020 | 4 | 16 | 32.85 | 34.95 | 280 |
| 44 | 19/07/2020 | 4 | 17 | 32.85 | 34.95 | 324 |
| 45 | 19/07/2020 | 5 | 19 | 32.85 | 34.95 | 150 |
| 46 | 19/07/2020 | 5 | 21 | 32.85 | 34.95 | 298 |
| 47 | 19/07/2020 | 5 | 22 | 32.85 | 34.95 | 296 |
| 48 | 19/07/2020 | 6 | 24 | 32.85 | 34.96 | 297 |
| 49 | 19/07/2020 | 6 | 25 | 32.85 | 34.96 | 269 |
| 50 | 19/07/2020 | 6 | 26 | 32.85 | 34.96 | 289 |
| 51 | 29/06/2021 | 1 | 95 | 31.68 | 34.55 | 108 |
| 52 | 29/06/2021 | 1 | 105 | 31.70 | 34.55 | 291 |
| 53 | 29/06/2021 | 2 | 965 | 34.56 | 31.70 | 140 |
| 54 | 07/07/2021 | 1 | 17 | 34.74 | 32.05 | 256 |
| 55 | 04/07/2022 | 2 | 1 | 32.35 | 34.84 | 327 |
| 56 | 04/07/2022 | 3 | 1 | 32.35 | 34.84 | 94 |
| 57 | 04/07/2022 | 4 | 2 | 32.36 | 34.84 | 151 |
| 58 | 04/07/2022 | 4 | 3 | 32.36 | 34.84 | 180 |
| 59 | 04/07/2022 | 4 | 4 | 32.36 | 34.84 | 29 |

**Movie S1** An example drone-based video of aggregated jellyfish. The video was taken at 7:45 AM on July 6, 2020. The video down-sampled from 30 fps to 2 fps, and shown sped up x10. https://www.youtube.com/watch?v=461N_J5ViY4

**Movie S2** An example drone-based video of the trajectories of aggregated jellyfish. Shows the overlaid trajectories of tracked jellyfish (green to red, corresponding with the time in the video) and passive drifters (white) overlaid on the same video shown in Movie S1. The discrepancy between the directionality of the jellyfish and the passive drifters exemplified here is representative of all the videos examined in this study. The video was taken at 7:45 AM on July 6, 2020. The video down-sampled from 30 fps to 2 fps, and shown sped up x10. https://www.youtube.com/watch?v=zEUGtpO4joo

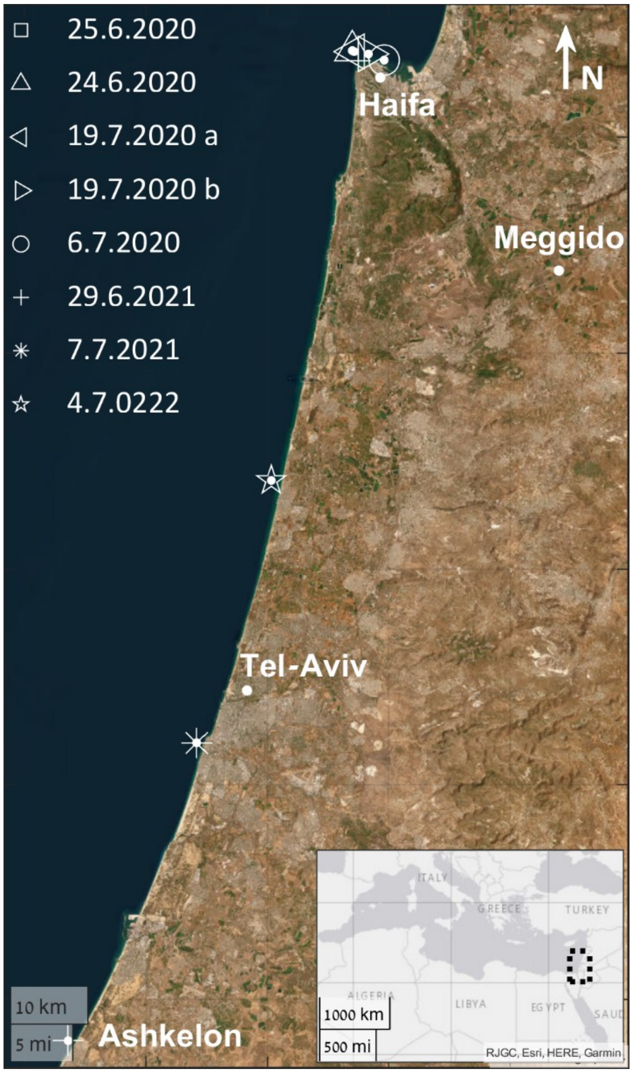


**Fig. S1** Locations of the field campaigns. Lower right panel shows the general location of the study area in the Southeastern Mediterranean Sea.

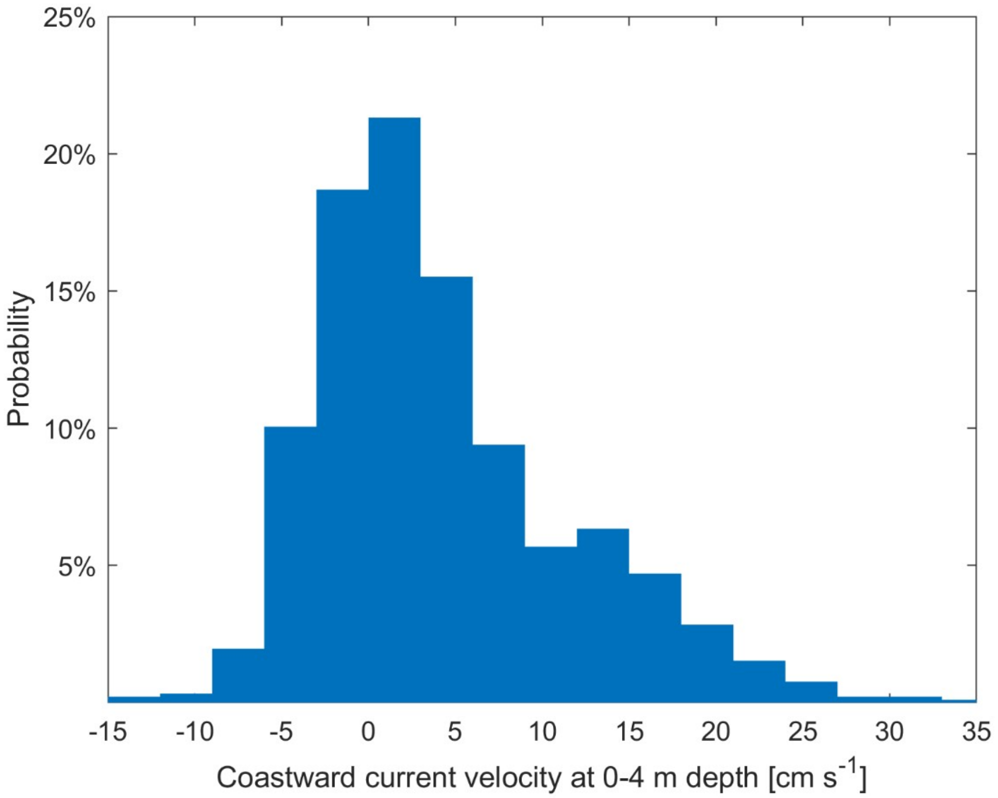


**Fig. S2** Summertime surface currents from long term ADCP measurements at a distance of 2 km from the coast. The data that is shown consists of measurements taken between 6AM and 10AM, during June and July of the years 2020 - 2022.

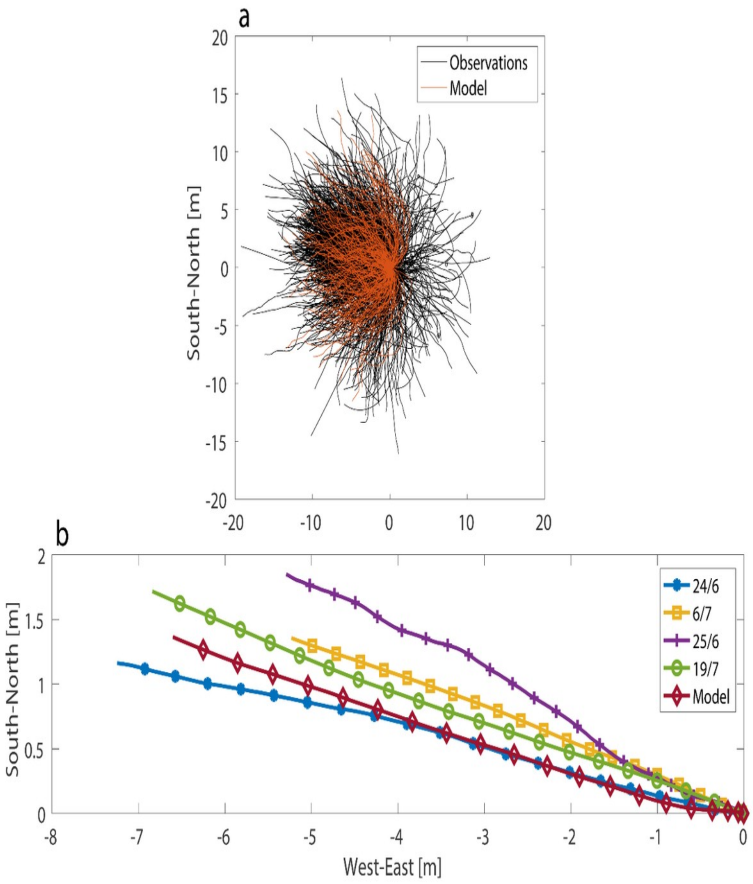


**Fig. S3** Jellyfish swimming trajectories. (a) Observed (black) and modeled (red) jellyfish active swimming trajectories in the east-west (x-axis) and north-south (y-axis) plane. (b) Modeled (red) and observed (blue, orange, purple, green) center of mass of active swimming trajectories. 24/6 and 6/7 were used by the GA, 25/6 and 19/7 were not.

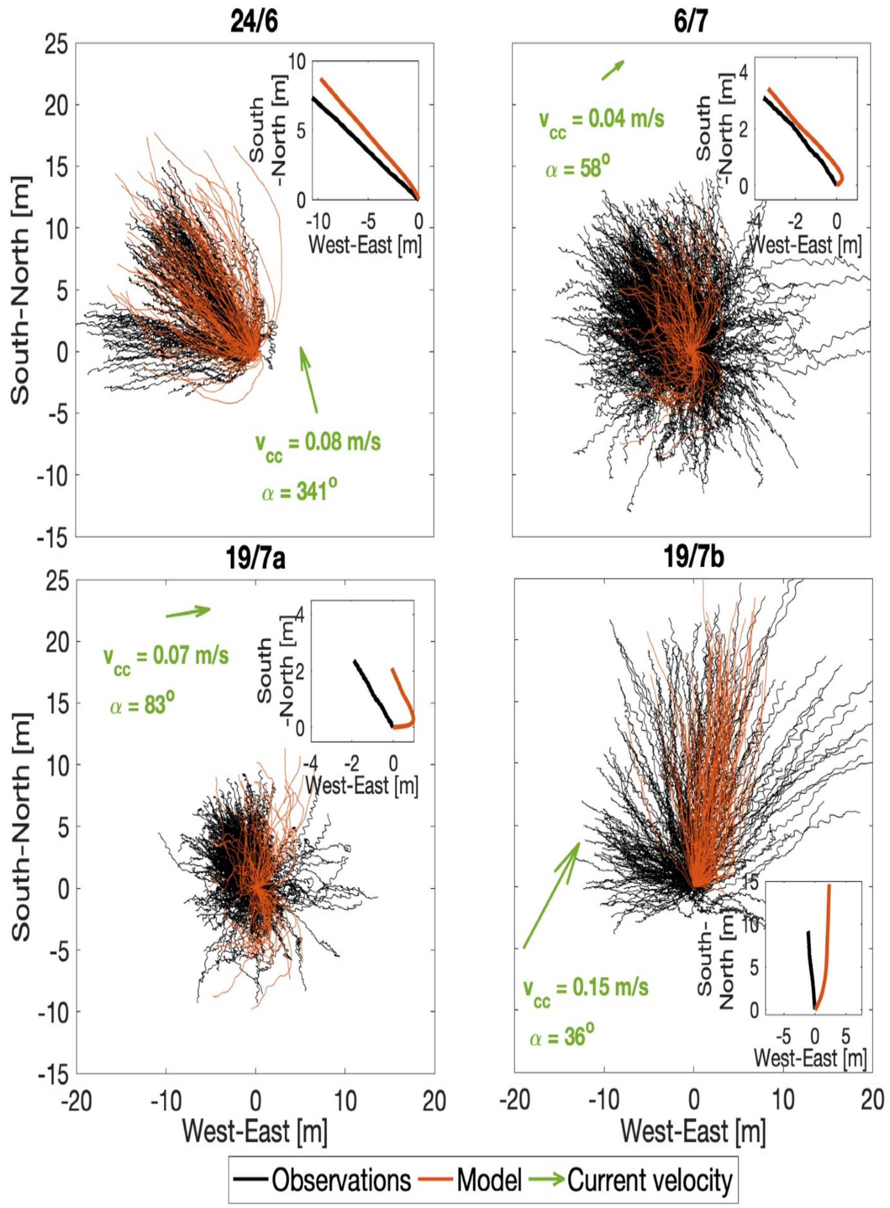


**Fig. S4** Observed (black) and simulated (red) trajectories under constant current ($v_{cc}$). The direction of the current for each observation is indicated by a green arrow within the figure. The small inner graph shows the center of mass of the observed (black) and modeled (red) trajectories in the same day. Note that the axis of the center of mass in the inner small graphs are not consistent in all plots.

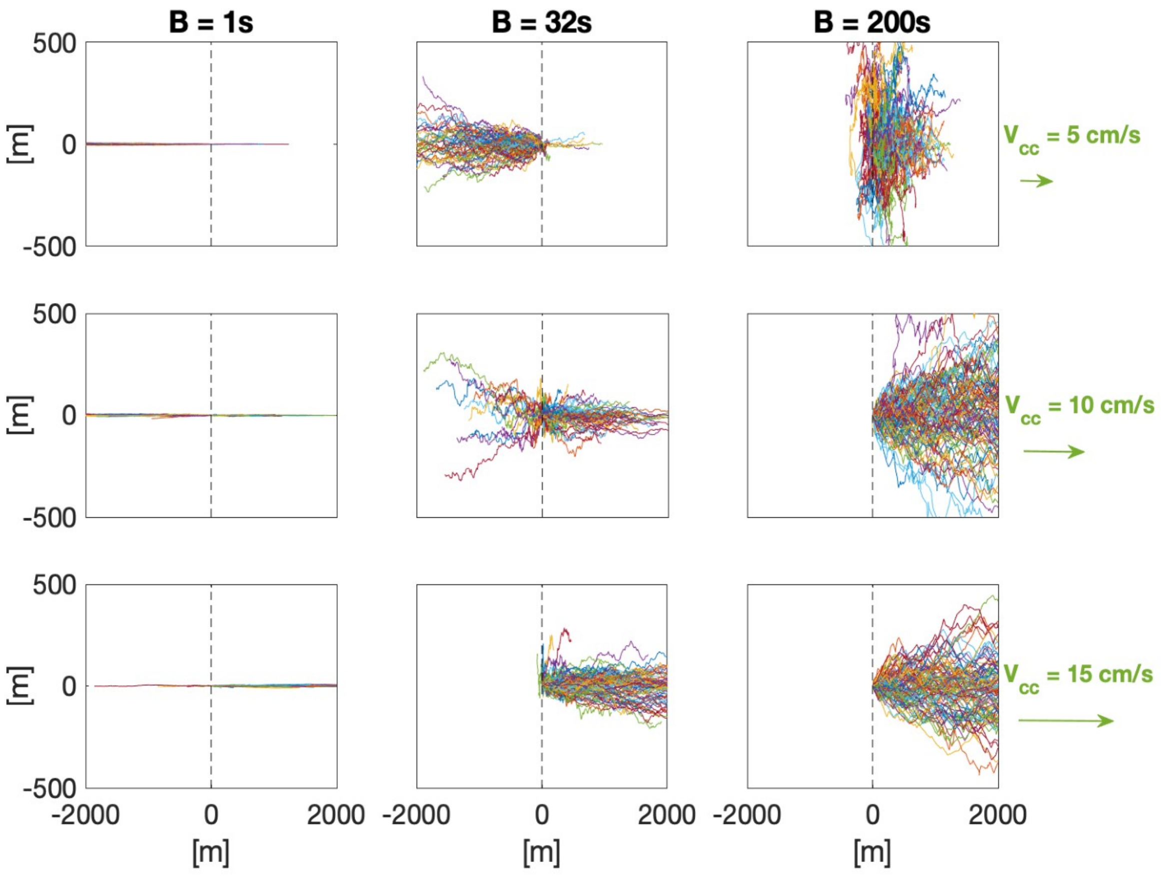


**Fig. S5** Jellyfish trajectories from the stranding simulations. Each row within the figure presents trajectories under the same constant current velocity ($v_{cc}$), while each column illustrates trajectories under the same reorientation time (B). Notably, there are discernible trends observed in the trajectories: As the constant current velocity ($v_{cc}$) increases, a greater number of jellyfish tend to propagate towards the east. As the reorientation time (B) increases, the span of the trajectories expands, and the directionality decreases. In such cases, jellyfish trajectories become more influenced by the passive current rather than their active swimming behavior.
